# ROS regulation of stigma papillae growth and maturation in *Arabidopsis thaliana*

**DOI:** 10.1101/2025.04.08.647846

**Authors:** Subramanian Sankaranarayanan, Sowmiya Devi Venkatesan, Thomas C. Davis, Sharon A. Kessler

## Abstract

Highly specialized stigma papillae cells play a critical role in plant reproduction. Their main purpose is to catch and interact with pollen, to mediate compatibility responses, to regulate pollen germination, and to guide pollen tubes to the transmitting tract so that the sperm cells carried in the pollen can be delivered to the female gametophyte to achieve double fertilization. In *Arabidopsis thaliana*, the stigma consists of single-celled stigma papillae that emerge from the apex of the fused carpels. Despite their critical function in plant reproduction, the molecular mechanisms that govern growth and maturation of stigma papillae remain poorly understood. Reactive Oxygen Species (ROS) have been implicated in stigma receptivity, but their roles in papillae development are less explored. Here we show that reactive oxygen species (ROS) also play different roles in stigma papillae development, with superoxide accumulating during the initiation and growth phase and hydrogen peroxide accumulating in mature papillae that are receptive to pollen. Reducing superoxide levels in the stigma by pharmacological treatments or over-expressing superoxide dismutase enzymes under an early stigma promoter inhibited stigma papillae growth, suggesting that ROS homeostasis is critical to papillae growth and differentiation for optimal pollination.

**Statements and Declarations:** The authors declare no competing interests

## Introduction

The female gametophytes of angiosperms are buried under layers of sporophytic tissue that protect and nurture the developing seeds post-fertilization. However, sexual reproduction requires that the male and female gametes be in direct contact. As a consequence, angiosperms have evolved an extraordinary tissue system, the reproductive tract, through which the male-derived pollen carries the immobile sperm through carpel tissue to the egg and central cell of the female gametophyte (Johnson *et al*., 2019; Ogawa and Kessler, 2023). The reproductive tract is a collection of highly specialized tissues, including the stigma, style and the transmitting tract, which collectively nurture and guide the pollen tube from its initial contact with the stigma to its final attraction to the ovule, enabling double fertilization (Crawford and Yanofsky, 2008). As the first point of contact for pollen, the stigma plays multiple critical roles in the sexual reproductive process, including catching and adhering pollen grains, assessing pollen compatibility, initiating pollen tube growth, and guiding the pollen tube towards the transmitting tract (Edlund *et al*., 2017; Heslop-Harrison, 1978; Heslop-Harrison and Shivanna, 1977; Takayama and Isogai, 2005; Wolters-Arts *et al*., 1998).

In Arabidopsis, the gynoecium (or pistil) consists of two carpels that initiate as a congenitally fused ring in the fourth floral whorl. As the primordial ring grows, the interior faces of the ring undergo postgenital fusion to form an elongated organ with two locules that will contain the ovules. After this fusion, the style is formed at the top of the pistil and gives rise to the stigma (Gasser and Robinson-Beers, 1993; Smyth *et al*., 1990). The stigma is a unique structure made up of highly specialized papillae that project outward, forming a a dome-shaped surface optimized for pollen capture. Arabidopsis papillae are single-celled and bell-shaped, with an elongated tip and a broader base. Auxin signaling and transcription factors such as NGATHA, HECATE, and STYLISH are required for specifying apical development to ensure that papillae form at the top of the pistil (Alvarez *et al*., 2009; Ballester *et al*., 2021; Gremski *et al*., 2007; Schuster *et al*., 2015). However, little is known about the molecular mechanisms contributing to papillae cell growth and differentiation after they are initiated.

Reactive oxygen species (ROS) function as important signaling molecules that regulate various aspects of plant growth and development (Mhamdi and Van Breusegem, 2018; Sankaranarayanan *et al*., 2020). For example, ROS regulate rapid cell expansion during morphogenesis in roots and leaves through effects on cell wall remodeling (Carol and Dolan, 2006; Schmidt *et al*., 2016). ROS accumulation in the active growth site correlates with the determination of cell shape of root hair cells (Takeda *et al*., 2008). In flowers, ROS mediates conical cell shaping in Arabidopsis petals through the modulation of microtubule ordering (Dang *et al*., 2018). While ROS have not been directly linked to cell growth and shape in the pistil, they are known to accumulate in stigmas, pollen tubes, and female gametophytes, where they play essential roles in double fertilization and seed production (Ali and Muday, 2024; Lan *et al*., 2017; Liu *et al*., 2021; McInnis *et al*., 2006a; Sankaranarayanan *et al*., 2020).

Stigma development in Arabidopsis consists of two stages: in the first stage, stigma papillae initiate at the apex of the fused carpels and grow out from the carpel surface. In the second stage, the papillae mature and become receptive to pollen while retaining the capacity for further growth in the absence of pollen. Given the importance of ROS in mediating growth process of various plant cell types and its functional role in pollination, we hypothesized that ROS also play roles in earlier stages of stigma papillae growth and maturation. Our findings reveal that superoxide accumulates in papillae during early stigma development when papillae are actively growing but not yet receptive to pollen. Superoxide accumulation drops at floral development stage 12 (when stigmas become receptive to pollen) before reappearing again in a second growth phase. Conversely, hydrogen peroxide levels are low during early growth stages but accumulates as the papillae mature. Pharmacological and genetic manipulation of superoxide accumulation reveal that superoxide accumulation promotes papillae growth during stigma development.

## Results and Discussion

### Superoxide accumulates in the stigma and displays distinct spatiotemporal patterns during stigma papillae growth

Previous studies using non-specific ROS detection showed that ROS accumulate in both immature and mature stigmas in many species, including Arabidopsis (Zafra *et al*., 2016). In plants, the main ROS that have been implicated in growth and development are superoxide and hydrogen peroxide (Martin *et al*., 2022). In order to determine if ROS has a potential role in stigma papillae growth and development, we first visualized the spatiotemporal patterns of superoxide and hydrogen peroxide accumulation in stigmas of wild-type (Col-0) Arabidopsis at different stages of development (stage 9 to stage 13) by staining pistils with the cell permeable fluorescent dyes PeroxyOrange 1 (PO1) to detect hydrogen peroxide and Dihydroethidium (DHE) to detect superoxide respectively (Chapman and Muday, 2021; Martin *et al*., 2022; Watkins *et al*., 2017) (Fig. 1). When stigmas from single inflorescences were stained and imaged from floral stages 9 through 15, clear differences were seen between the timing of hydrogen peroxide and superoxide accumulation in stigma papillae. DHE detected superoxide in stage 9 stigmas near the site of stigma papillae initiation and in growing papillae up until stage 11/early stage 12, coinciding with the first anisotropic growth phase (Fig. 1C-D). No superoxide was detected by DHE staining in stage 12b-13 stigmas, coinciding with when papillae start to become receptive to pollen. After stage 13, superoxide was only detected in pollen and not in papillae in the open flowers of the inflorescence or in unpollinated stage 14 flowers (Fig. 1D and 2A-D). PO1 staining of hydrogen peroxide showed an opposite trend to DHE staining. Hydrogen peroxide was not detected when superoxide was present in floral stages 9 through 11 and started to appear by stage 12b. In stage 13 and 14 stigmas from open flowers, hydrogen peroxide was not detected in papillae, but was present in germinated pollen on stigmas (Fig. 1E-F and 2H)). However, in stage 14 stigmas from emasculated flowers, hydrogen peroxide levels remained high in papillae cells (Fig. 2G). These data suggest that superoxide accumulation may be associated with stigma papillae growth while hydrogen peroxide could have a role in stigma receptivity since it accumulates when the stigma reaches maturity and is downregulated after pollination.

**Figure 1.**
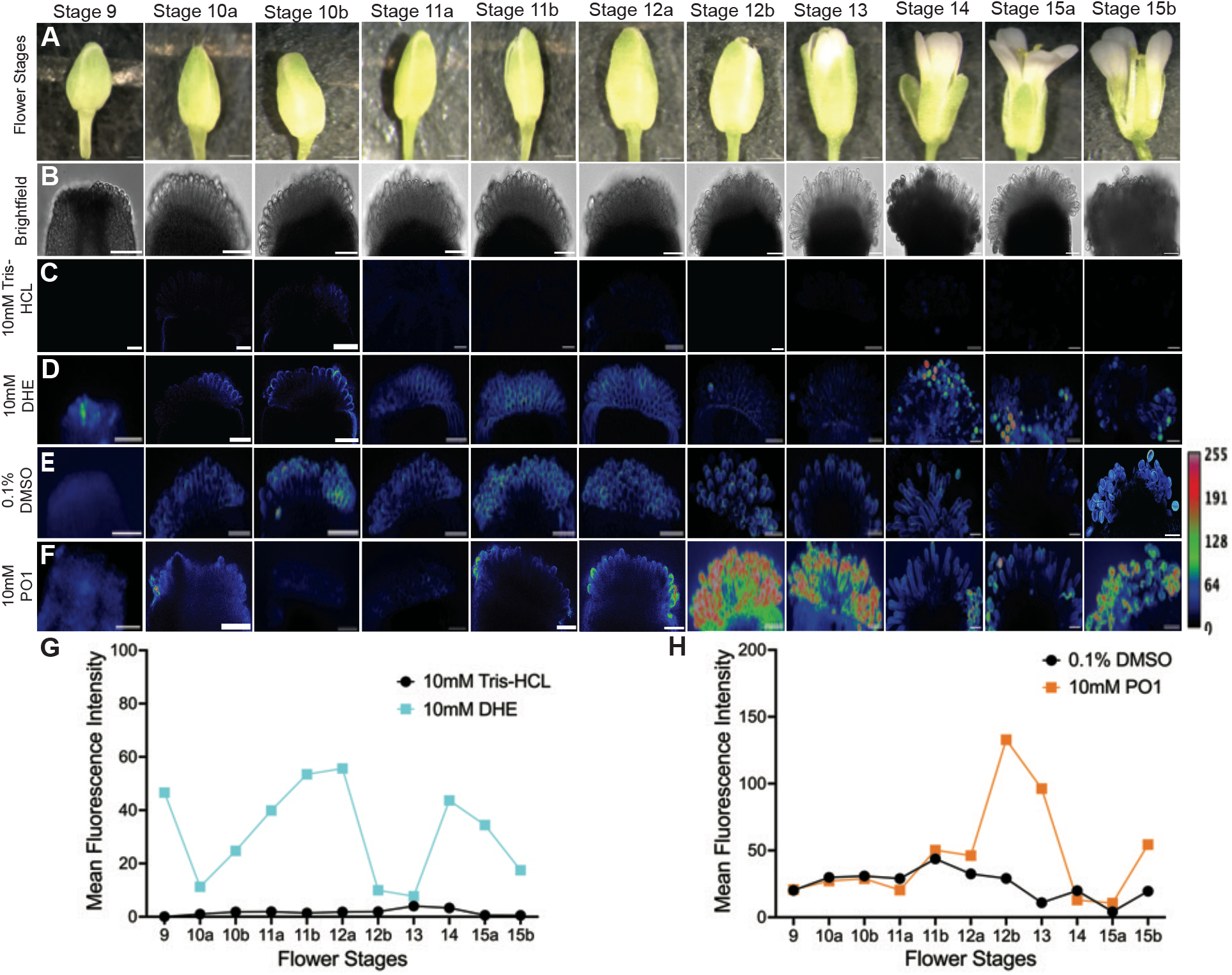
Intracellular superoxide levels are high in papillae in the early growth phase while hydrogen peroxide accumulates at the transition to maturity. Superoxide and hydrogen peroxide detection in stigma developmental stages using Dihydroethidium (DHE) and Peroxy Orange 1 (PO1), respectively. (A) Representative images of flowers from stage 9 to stage 15b. Scale bar = 1 mm. (B) Representative brightfield images of stigma from stage 9 to stage 15b used for imaging. Scale bar = 50 µm. (C-F) Maximum intensity projections of stigmas under different treatments, mock treatment 10mM Tris-HCL (C) and 10 mM DHE (D); mock treatment 0.1 % DMSO (E) and 10 mM PO1 (F). Scale Bar = 50 µm. (G, H) The plots represent the mean fluorescence intensity of images under mock and stained conditions for DHE and PO1, respectively. Stigmas in each treatment shown in the figure were collected from a single inflorescence and are representative of three experimental replicates.

**Figure 2.**
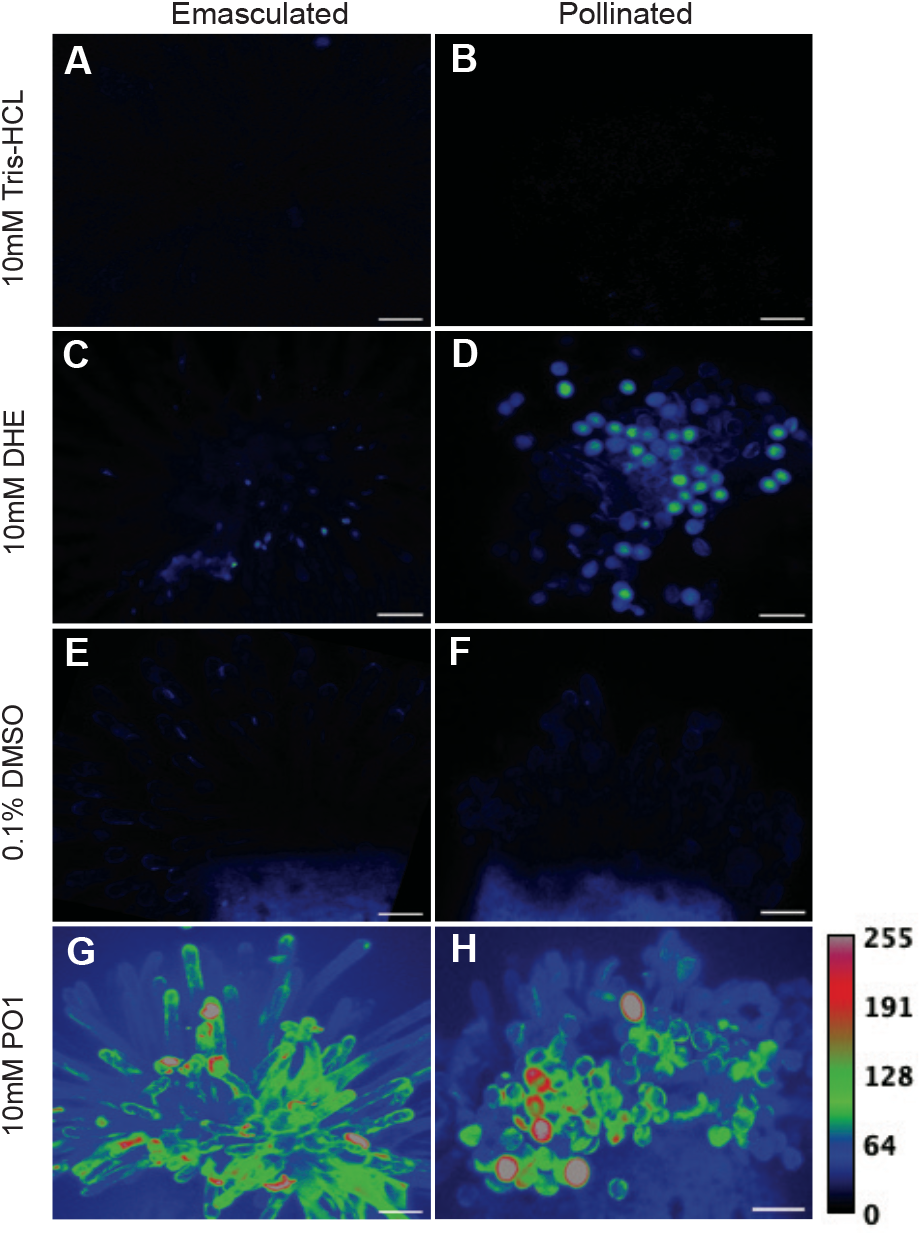
Intracellular hydrogen peroxide levels decrease in stigma papillae after pollination, while both superoxide and hydrogen peroxide are present in adhered pollen. Superoxide and hydrogen peroxide detection in emasculated and pollinated pistils using DHE and PO1, respectively. (A, C, E, G) Maximum intensity projections of unpollinated stigmas (two days after emasculation) under different treatments, control 10mM Tris-HCL (A) and 10 mM DHE (C); control 0.1 % DMSO (E) and 10 mM PO1 (G). (B, D, F, H) Maximum intensity projections of pollinated stigmas (Stage 15a) under different treatments, control 10mM Tris-HCL (B) and 10 mM DHE (D); control 0.1 % DMSO (F) and 10 mM PO1 (H). Scale Bar = 50 µm.

PO1 and DHE are both cell permeable and therefore react with intracellular ROS. The colorimetric ROS stains, Nitrotetrazolium blue (NBT) and 3,3’-Diaminobenzidine (DAB) are non-targeted and can detect both intracellular and extracellular superoxide and hydrogen peroxide, respectively (Ben Rejeb *et al*., 2015; Brandes and Janiszewski, 2005; Martin *et al*., 2022). Superoxide accumulation patterns revealed with NBT staining were similar to the DHE results in stage 9-12 stigmas. Superoxide was detected at the top of the pistil before papillae initiation and continued through floral stage 11. NBT staining was reduced significantly in stage 12 stigmas. In contrast to the DHE results, NBT staining reappeared in stage 13 when the stigma is fully mature, yet papillae are still growing in unpollinated stigmas (Fig. 3A). DAB staining (Fig. 3B) gave similar results to the PO1 detection of hydrogen peroxide in early stages of stigma development. DAB staining was very weak when NBT staining was high in stages 9 and 10. In contrast to PO1 staining, DAB staining increased in intensity slightly earlier at stage 11, which could reflect extracellular accumulation of hydrogen peroxide, although the resolution of DAB staining is not sufficient to make strong conclusions about subcellular localization. Unpollinated stage 12 and 13 stigmas had stronger DAB staining compared to the early stages, similar to the results with PO1 (Fig. 3B and 2G).

**Figure 3.**
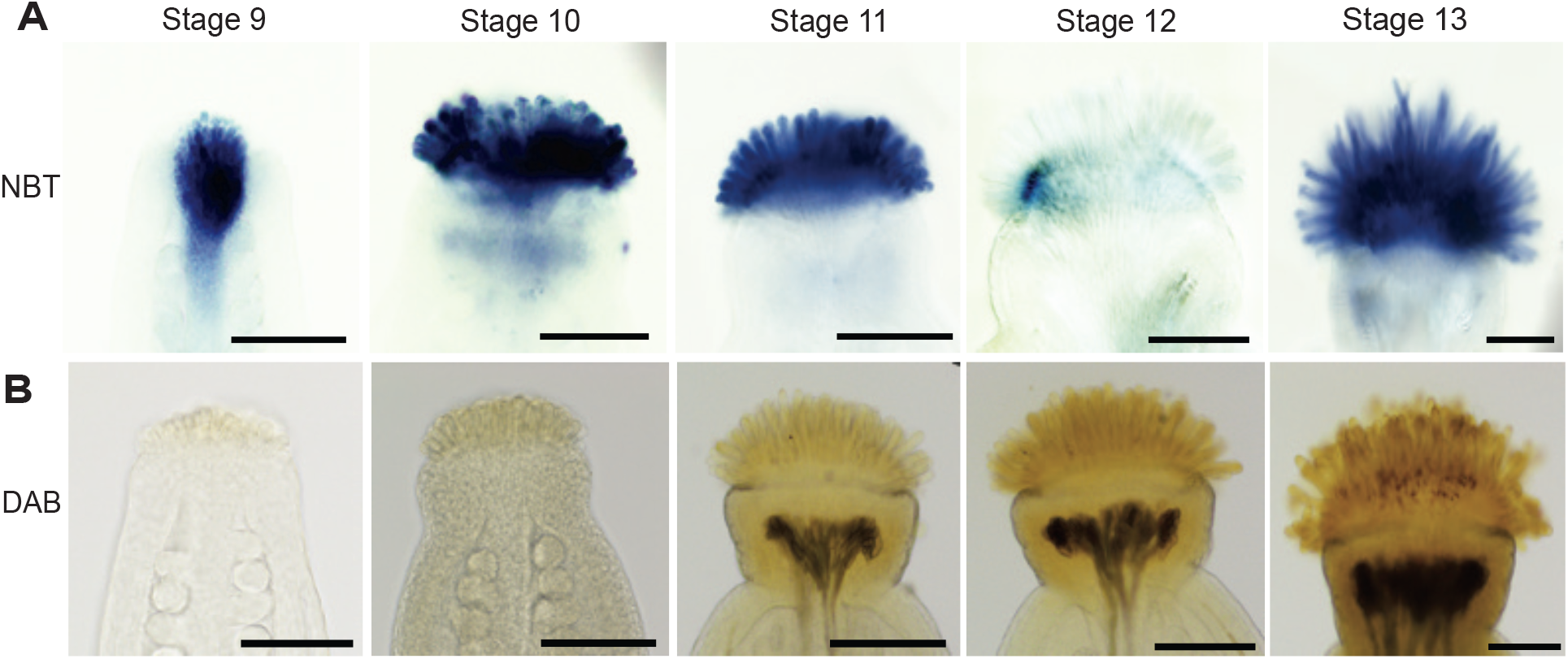
Colorimetric ROS detectors reveal that superoxide levels are high in papillae in the early growth phase while hydrogen peroxide accumulates at the transition to maturity. (A) NBT detection of superoxide in stage 9-13 Col-0 stigmas. (B) DAB staining of hydrogen peroxide in stage 9-13 Col-0 stigmas. Scale bars = 100 µm

### Superoxide accumulation during stigma development is correlated with papillae growth

DHE and NBT staining suggested that superoxide may be important for papillae cell growth. To test this hypothesis, we used pharmacological and genetic approaches to manipulate superoxide levels in developing stigmas. We used Diphenyleneiodonium Chloride (DPI) to inhibit NAD(P)H oxidase-mediated superoxide production at the plasma membrane (extracellular superoxide) and the superoxide scavenger *n*-propyl gallate (*n*PG) to manipulate superoxide accumulation in developing papillae (Lambert *et al*., 2008; Reddan *et al*., 2003). Pistil-feeding experiments with both DPI and nPG resulted in a significant reduction in stigma papillae length (Fig. 4). Treatment of stigmas with 1 µM DPI and 10 µM DPI resulted in a 22% and 32% decrease in papillae length respectively compared to control stigmas (Fig. 4C). Similarly, treatment of stigmas with 10 µM and 100 µM nPG resulted in a ∼35% decrease in papillae length compared to control stigma (Fig. 4D). In both treatments, NBT staining levels were significantly reduced compared to the control stigmas (Fig. S1 A-D), indicating that we had effectively reduced superoxide in the papillae.

**Figure 4.**
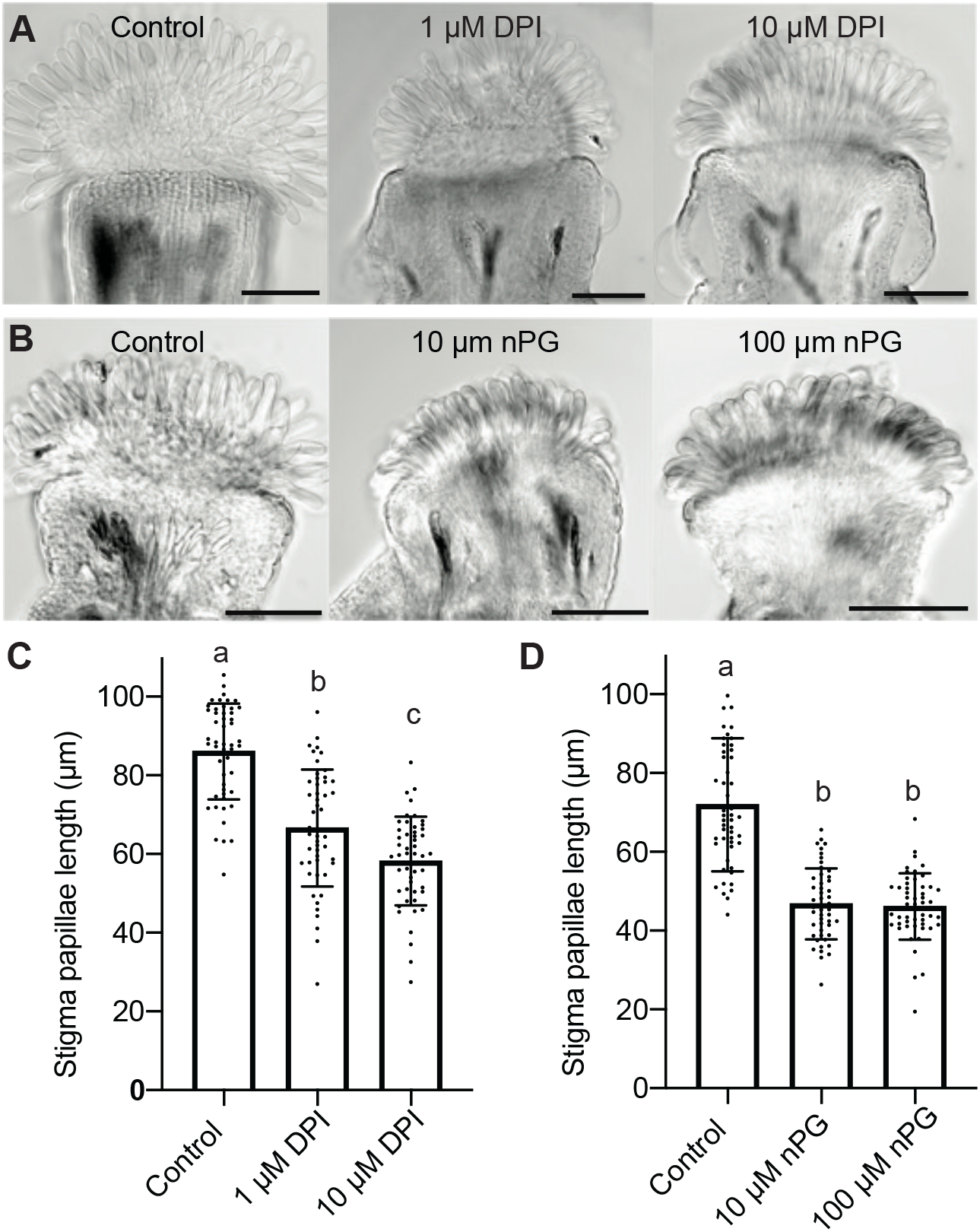
Reduction of superoxide levels through treatment with inhibitors or scavengers reduces stigma papillae growth. (A) Representative image of a control stigma vs Diphenyleneiodonium chloride (DPI) treated stigmas. (B) Representative image of a control stigma vs n-Propyl gallate (nPG) treated stigmas. (C) Average stigma papillae length of control vs DPI treated stigma (1µM and 10 µM) (n=50 per treatment). (D) Average stigma papillae length of control vs nPG treated stigmas. (n=50 per treatment). Error bars indicate ± SD. Significance was determined by a one-way ANOVA with Tukey test (letters indicate p≤0.05). Scale bars =100µm

To confirm that superoxide plays a role in stigma papillae growth, we aimed to use a genetic approach to reduce the level of intracellular superoxide during stigma development. We reasoned that overexpression of superoxide dismutase (SOD) enzymes in early stages of papillae development would decrease superoxide accumulation specifically in these cells as opposed to our pistil-feeding approach that could potentially affect other aspects of pistil development leading to papillae growth defects. The S-Locus Related 1 (SLR1) promoter from *Brassica olereracea* is commonly used in stigma research, but it is mainly active in later stages of stigma development in Arabidopsis (Dwyer *et al*., 1994). We therefore first needed to identify a promoter that is active in early stages of papillae development in order to genetically manipulate superoxide levels. We used promoter::GUS analysis of genes that are expressed highly in stigmas to determine that the promoter of a gene of unknown function, AT5g53710, referred to hereafter as CHLORIS after the Greek goddess of flowers, is active in stage 9-13 papillae (Fig. S2 A-C) and is therefore sufficient for ectopically expressing SOD genes in early stigma development.

SODs are enzymes that scavenge superoxide (O_2_^-^) radicals. They have been shown to be involved in antioxidant-mediated defense in plants (Kliebenstein *et al*., 1998; Martin *et al*., 2013). The Arabidopsis genome encodes seven SODs that are classified by their active site metal co-factors (manganese, copper, zinc, or iron). SODs are localized in a variety of subcellular compartments. In Arabidopsis, MnSOD (MSD1) is localized in mitochondria, while the three *CuZnSODs* (*CSD1, CSD2, CSD3*) are localized in the cytosol, peroxisomes, and chloroplasts, respectively. The three *FeSODs* (*FSD1, FSD2 and FSD3*) are localized in chloroplasts and peroxisomes (Kliebenstein et al., 1998). We hypothesized that overexpressing SODs that localize in mitochondria and the cytotosol would be most likely to affect superoxide levels in developing papillae.

We first expressed the mitochondria-localized *MSD1* under the CHLORIS promoter in wild-type stigmas. We identified three independent transformants (lines L3, L4 and L7) that accumulated 3.5 to 4-fold *MSD1* mRNA in their stigmas compared to the wild-type control (Fig. S3A). Stigma papillae length was reduced in all three MSD1-OX lines compared to non-transgenic stage 13 mature flowers (Fig. 5A, B). At stage 13, Line 3 and 4 stigmas had almost 25% reduction in papillae length compared to the control, while Line 7 showed a 34% reduction in papillae length compared to the control (Fig. 5B). There were no noticeable differences in other aspects of floral morphology between the transgenic *MSD1-OX* lines and the control (Fig. S3B). We selected Line 7 for stage-specific analysis of papillae growth since it had the strongest effect by floral stage 13. We first estimated superoxide levels in the stigma of the MSD1-OX line at different stages of development through NBT staining. We observed a significant reduction in NBT staining in stigmas of the *MSD1-OX* line at stages 9-11 and 13 in comparison to the wild-type control (Fig. 4D and S4C). Stage 12 stigmas had little to no NBT staining in both the MSD1-OX line and the Col-0 control, as expected based on our previous experiments (Fig. 3A). When papillae lengths were compared over the course of stigma development, MSD1-OX papillae were significantly shorter than wild-type papillae at all stages examined (Fig. 5C). A similar result was found when a cytosol-localized superoxide dismutase, CSD1/SOD1, was overexpressed in developing papillae. In three independent pCHLORIS::CSD1 lines, papillae were significantly shorter than wild-type controls (Fig. 6 A, B) and NBT staining intensity was also greatly reduced (Fig. 6C). Taken together, the results from our genetic study overexpressing superoxide dismutase enzymes support a role for intracellular superoxide in the regulation of stigma papillae growth.

**Figure 5.**
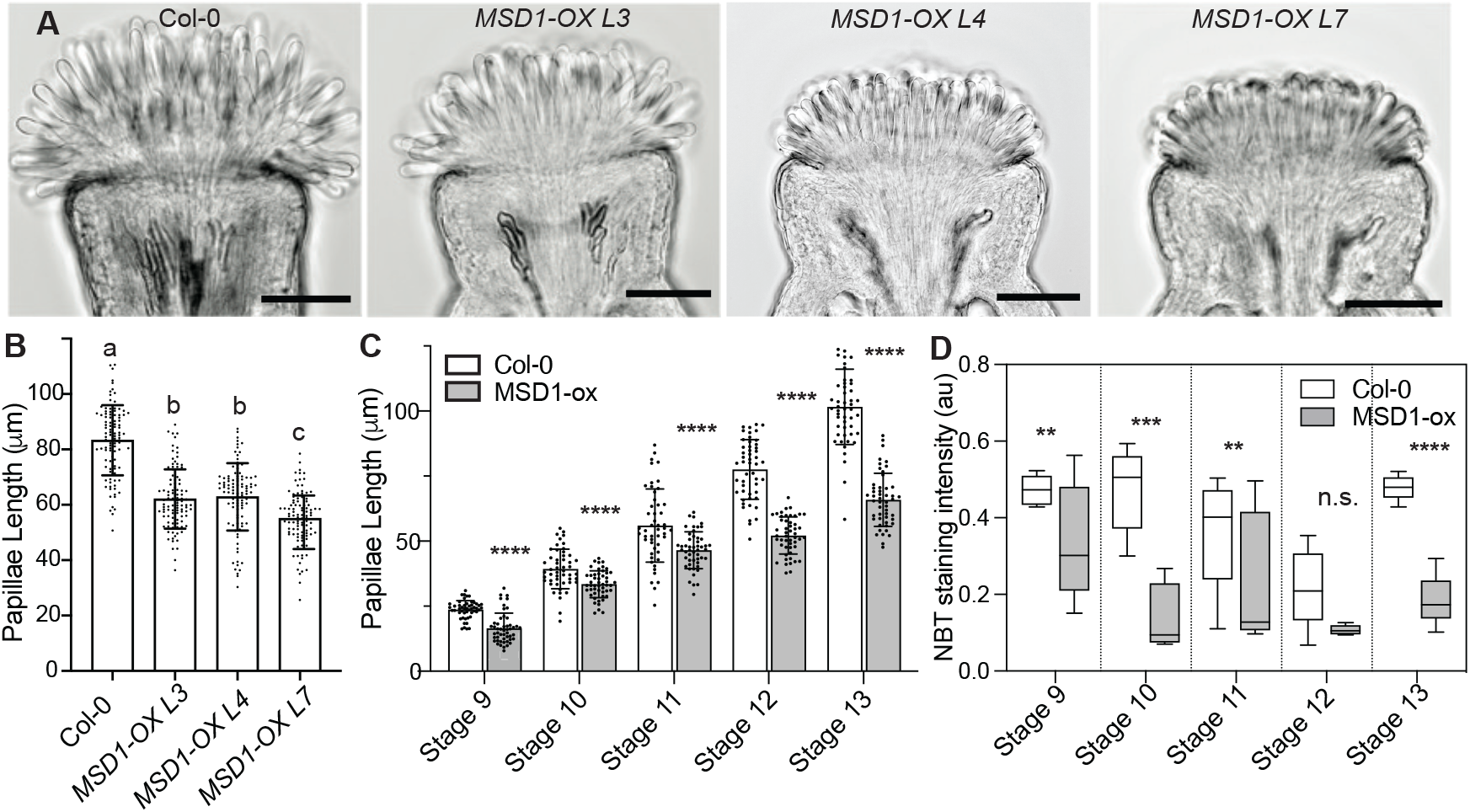
Overexpression of *MSD1* in developing stigmas reduced superoxide levels and suppresses papillae growth. (A) Representative images of stage 13 wild-type stigma and stigmas of three independent transgenic stigma specific *MSD1* overexpression lines (L3, L4, L7) in wild-type background. Scale bars=100 µm. (B) Average stigma papillae length of stage 13 wild-type vs three *MSD1* overexpression lines (n=100 per genotype). Significance was determined by a one-way ANOVA with Tukey test (letters indicate p≤0.05). (C) Average papillae length of *Col-0* vs *MSD1-OX* line (L7) at different stages of floral development (n=50 papillae per stage). (D) Average NBT stain intensity in stigmas from different developmental stages of *Col-0* and *MSD1-OX* line 7. (n=5) Error bars indicate ± SD. Asterisks in C-D indicate significant differences (*P<0.05, **P<0.01, ***P<0.001, ****P<0.0001) determined by ordinary One-way ANOVA with Dunnett’s test.

**Figure 6.**
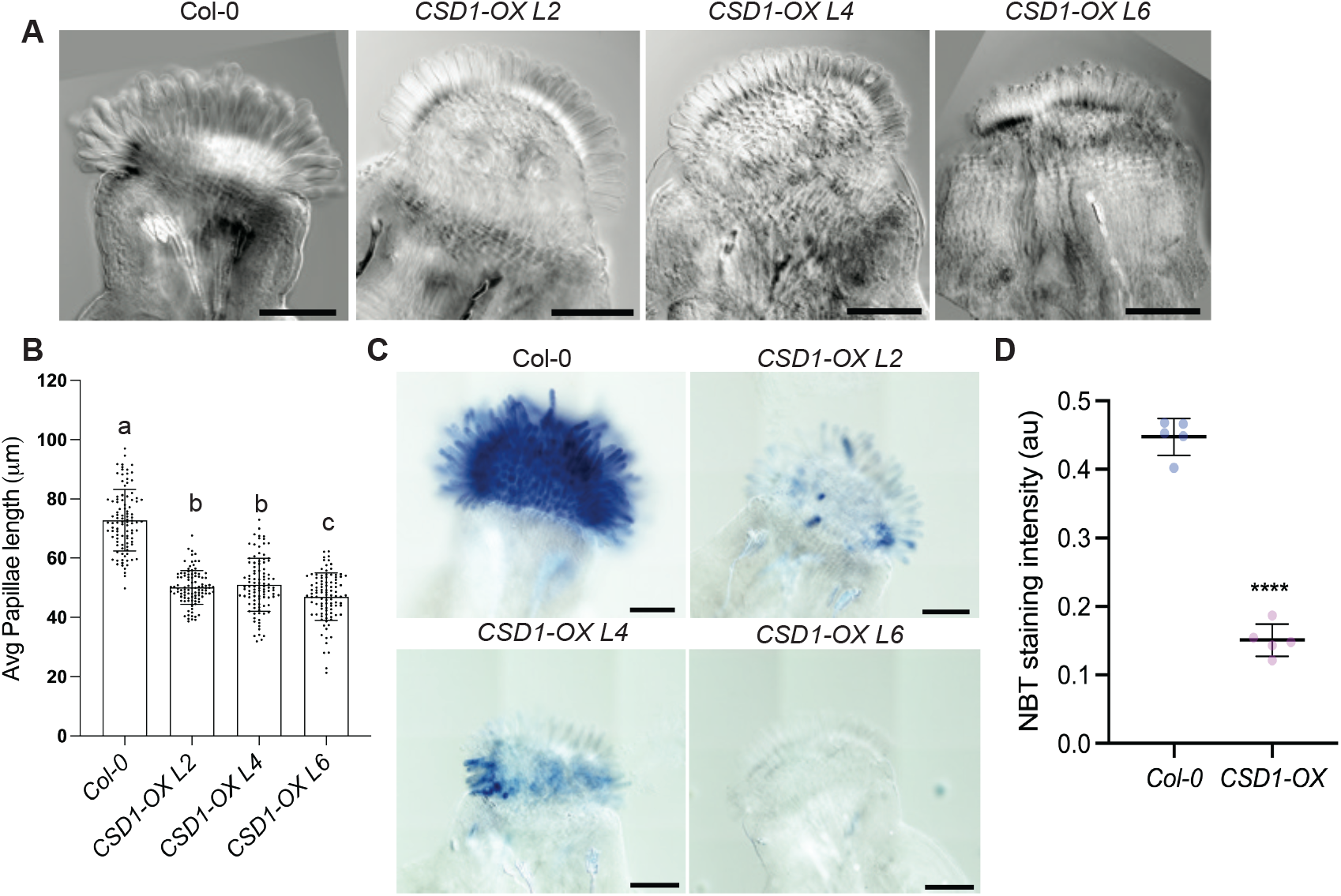
Overexpression of *CSD1* in developing stigmas reduces superoxide levels and suppresses papillae growth. (A) Representative images of stage 13 wild-type stigma and stigmas of three independent transgenic stigma specific *CSD1* overexpression lines (L3, L4, L7) in a Col-0 background. Scale bars=100µm. (B) Average stigma papillae length of stage 13 wild-type vs three *CSD1* overexpression lines (n=100 per genotype). Significance was determined by a one-way ANOVA with Tukey test (letters indicate p≤0.05). (C) CSD1-OX lines have reduced NBT staining in stage 13 stigmas. (D) Quantification of NBT staining in CSD1-OX.

### A potential link between superoxide and microtubules during papillae growth

The cytoskeleton plays a major role of controlling growth direction of the cells by guiding the delivery of Golgi-derived vesicles and the cellulose synthase machinery (Geisler *et al*., 2008). Both microtubules and actin filaments are sensitive to ROS (Lee *et al*., 2017; Livanos *et al*., 2012). It is possible that cytoskeleton dynamics during the growth of stigma papillae are modulated by changes in stigmatic ROS levels. Therefore, we tested whether DPI and nPG treatments affect microtubule arrangements in developing stigma papillae. In control papillae, microtubules were arranged transversely in papillae, consistent with anisotropic growth (Fig. 7). However, in both ROS inhibitor treatments, transverse microtubule arrays were not present (Fig. 7), indicating depolymerization of microtubules in response to perturbed superoxide accumulation in stigma papillae.

**Figure 7:**
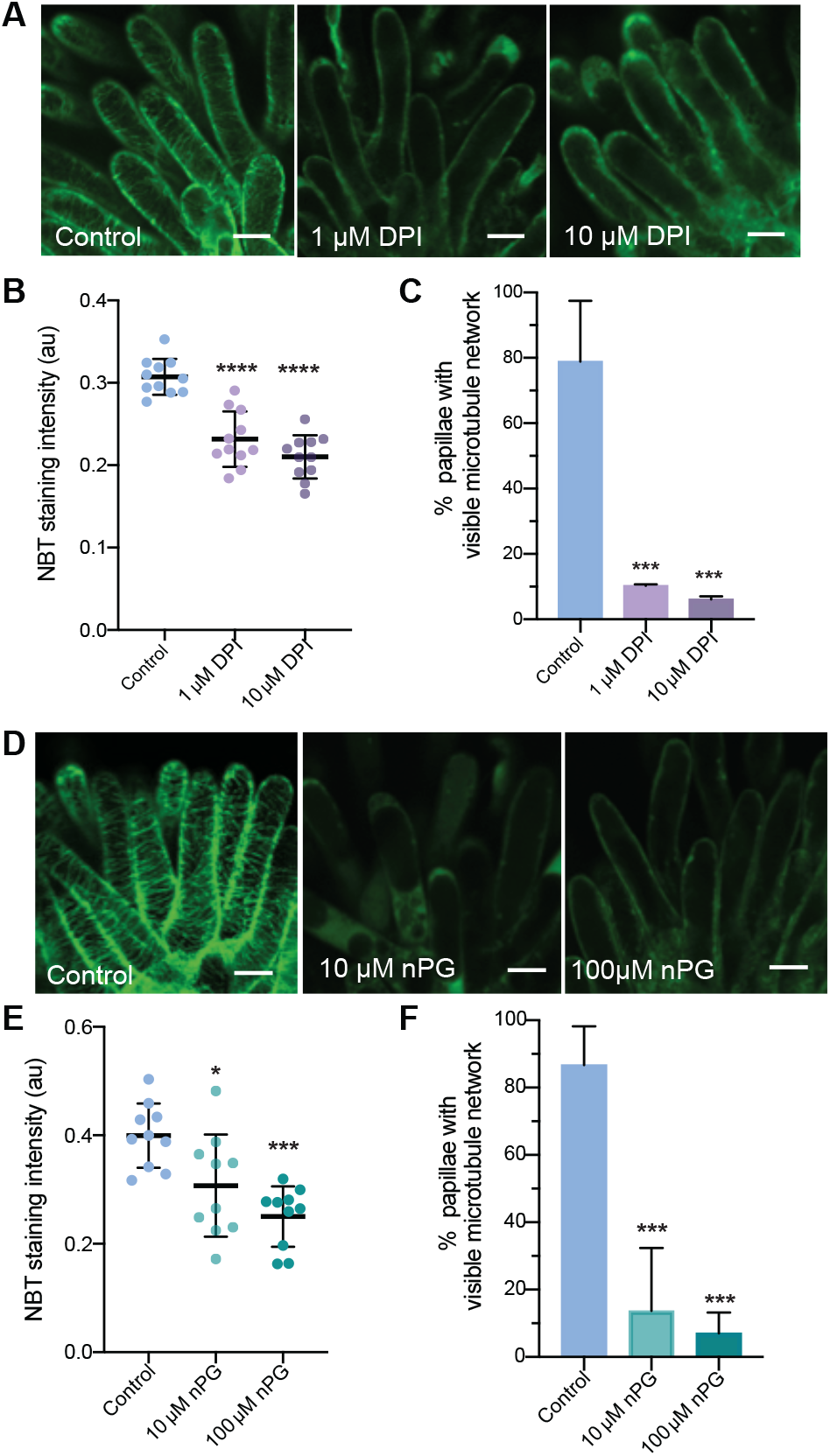
Pharmacological reduction of superoxide disrupts microtubule organization in papillae. (A) Representative confocal images of papillae expressing MAP65.1-citrine in stage 13 Col-0 control vs pistils treated with superoxide inhibitor DPI (1 µM and 10 µM). (B) Quantification of superoxide levels in DPI-treated stigmas by NBT staining. (C) Percentage of stigma papillae with visible microtubule network from control and following nPG treatments (n= at least 50 papillae for each treatment). (D) Representative confocal images from nPG (10 µM and 100 µM) treated stage 13 stigmas. (E) Quantification of superoxide levels in nPG-treated stigmas. (F) Percentage of stigma papillae with visible microtubule network from control and following DPI treatments (n= at least 50 papillae for each treatment). Error bars indicate ± SD. Asterisks indicate significant differences (****P<0.0001, ***P<0.001, *P<0.05) determined by ordinary One-way ANOVA with Dunnett’s test. Scale bars = 5 µM.

## Discussion

Stigma papillae play a critical role in facilitating pollination and optimal seed production. Our study revealed that ROS play an important role in ensuring that papillae grow and become receptive to compatible pollen. Arabidopsis stigma papillae achieve their elongated shapes via a polarized anisotropic diffuse growth mechanism that occurs in two phases. In the first phase, papillae initiate and increase in diameter and length while also accumulating superoxide (Fig. 1-3). The end of phase I coincides with a decrease in superoxide accumulation and an increase in hydrogen peroxide accumulation that is linked to stigma receptivity that starts in stage 12 of floral development. In phase 2 unpollinated stigmas, papillae continue to elongate while also accumulating hydrogen peroxide. In contrast, pollination leads to a reduction in hydrogen peroxide in papillae cells and a cessation of growth (Fig. 2). Pharmacological and genetic manipulation of superoxide levels in developing papillae revealed that superoxide accumulation promotes papillae growth (Fig. 4-5).

In plants, cell size and shape are regulated by localized extension of the cell wall leading to anisotropic expansion driven by turgor (Smith and Oppenheimer, 2005). Cell expansion requires both cell wall loosening and deposition of new cell wall (reviewed in (Bashline *et al*., 2014)). Cell wall rigidity depends on cellulose, the major component of the cell wall, and the extent to which it crosslinks non-cellulosic polysaccharides such as hemicellulose and pectin (Cosgrove, 2005). ROS have been shown to be involved in regulating cell expansion through cell wall modification (Schmidt *et al*., 2016). Extracellular superoxide can be converted into H_2_O_2_ through the activity of superoxide dismutase, after which it can aid in cross-linking the cell wall or can be converted into hydroxyl radicals (.OH) which causes cell wall loosening (Richards *et al*., 2015). In the cytosol, ROS have also been shown to affect microtubule dynamics, which in turn can affect cellulose deposition (Livanos *et al*., 2012; Schmidt *et al*., 2016). In Arabidopsis root tips, low ROS levels interfered with microtubule (MT) dynamics leading to MT instability in the NADPH oxidase mutant *root hair defective 2 (rhd2)* and in DPI-treated root tip cells (Livanos *et al*., 2012). In our experiments, superoxide accumulated in early stages of papillae initiation and growth, where cell wall loosening and microtubule-guided cellulose synthesis would be necessary for rapid expansion. Disruption of both intracellular and extracellular superoxide accumulation led to impaired papillae elongation in our experiments with ROS inhibitors/scavengers and with overexpression of superoxide dismutases (Fig. 4-6). Both DPI and nPG treatments disrupted microtubules in papillae (Fig. 7), indicating that maintenance of optimal intracellular and extracellular ROS levels may be important for regulating cytoskeletal dynamics. Whether this is a direct or indirect effect of ROS and its connections to cell metabolism remains to be determined.

While our results suggest a role for superoxide in stigma papillae initiation and expansion during early development, other studies have implicated ROS in stigma receptivity to pollen in a variety of species. DCFH_2_-DA, a fluorescent stain that detects both hydrogen peroxide and superoxide, detected ROS in immature and mature stigmas of other species with dry stigmas covered in a thin proteinaceous layer known as the pellicle as well as in species with wet stigmas containing exudates rich in lipids and/or carbohydrates (Hiscock *et al*., 2007; McInnis *et al*., 2006a; Zafra *et al*., 2016). Pre-treatment of mature *Senecio squalidas* and Arabidopsis stigmas with sodium pyruvate to scavenge hydrogen peroxide eliminated most of the DCFH_2_-DA staining, indicating that hydrogen peroxide makes up most of the ROS detected in mature *Senecio* and Arabidopsis stigmas (McInnis *et al*., 2006a). This result is consistent with our findings that hydrogen peroxide accumulates mainly after stage 12 in Arabidopsis stigma development. In addition to accumulating ROS, both wet and dry stigmas also accumulate peroxidases at maturity (Dupuis and Dumas, 1990; Galen and Plowright, 1987; McInnis *et al*., 2006b; Seymour and Blaylock, 2000). Peroxidase activity has been used as a marker for stigma receptivity to pollen in many species (Dafni and Maues, 1998; Galen and Plowright, 1987), however the link between peroxidase activity and ROS accumulation in stigmas is still not understood since peroxidases can both consume and produce hydrogen peroxide (Bolwell *et al*., 1999; Bolwell *et al*., 1995). One possibility is that peroxidases act to balance ROS levels in the stigma in order to promote interactions with arriving pollen grains while preventing cytotoxicity. Thus, maintenance of the proper balance of different ROS species is critical for both stigma papillae growth and function at maturity.

The sole function of stigma papillae is to interact with pollen. In compatible pollinations, communication with a papillae cell leads to rapid pollen adhesion and hydration and the growth of a pollen tube into the cell wall of the papillae cell in order to direct it towards the transmitting tract (Kandasamy *et al*., 1994). Our experiments showed that hydrogen peroxide levels are highest in mature stigmas that have not been pollinated. The receptor-kinase FERONIA regulates stigma ROS accumulation in both compatible and incompatible pollinations. Live imaging experiments showed that ROS play an important role in regulating pollen interaction with the stigma on short time scales (Liu *et al*., 2021; Zhang *et al*., 2021). Liu and co-authors showed that ROS levels detected by H2DCFA decrease within 1 minute of pollen interaction with a receptive papillae cell and that this reduction is necessary for pollen adhesion and hydration (Dwyer *et al*., 1994; Liu *et al*., 2021). These experiments were conducted with a general ROS stain that detect both superoxide and hydrogen peroxide, but our results suggest that the ROS involved is likely to be hydrogen peroxide.

### Conclusions and future directions

The interplay between different ROS molecules, papillae growth, and receptivity is obviously complex and must be balanced with the potential cytotoxicity of ROS. A similar scenario was observed in plant meristems where hydrogen peroxide accumulation coincides with decreased superoxide accumulation in plant stem cells and the resulting superoxide/hydrogen peroxide balance controls plant stem fate (Zeng *et al*., 2017). The next challenge in the stigma papillae model will be to determine whether a causative relationship exists between dynamic ROS accumulation in developing papillae and the cytoskeletal and cell wall changes that lead to directional growth and pollen receptivity.

## Materials and Methods

### Plant Growth and Materials

All plants used in this study were in the *Arabidopsis thaliana* Columbia-0 (Col-0) background. Seeds were sterilized in 30% sodium hypochlorite, 0.1 % Triton-X 100 for 10 minutes, followed by 3 washes in sterile water and a 2-3-minute incubation in 70% ethanol. Following sterilization, seeds were plated on ½ Murashige and Skoog (MS) media (Murashige and Skoog, 1962) with 0.6% phytoagar, stratified at 4 ºC for 2 days, and then moved to a light room. Seven-day-old seedlings were transplanted to soil and grown at 21-22 ºC in (16h:8h) long day conditions. Plants that had already produced at least 5 stage 14 or later flowers were used in the experiments.

### Floral Staging

Floral Staging was largely based on the stages defined by (Alvarez-Buylla *et al*., 2010), with some modifications since the original study was done with the Landsberg *erecta* (Ler) background while our study used a Col-0 background. In floral Stage 6 the gynoecium initiated in whorl 4. Stage 7 was identified by the appearance of anther filaments. Anther lobing was used to classify stage 8 flowers. Petal sizes of 45 to 200 µm were used as the stage 9 landmark. Stages 10 and 11 were identified by petals being longer than the short anthers but shorter than the long anthers (Alvarez-Buylla *et al*., 2010; Smyth *et al*., 1990). Style morphology was used to distinguish between stages 10 and 11. The style becomes more pronounced and style height surpassed stigma papillae length as flowers entered stage 11. Stage 12 was defined as the stage during which the anthers dehisced and petals were longer than the long stamens. After anthesis, flowers were classified as stage 13 flowers.

### Fluorescent staining and quantification of ROS in stigmas

H_2_O_2_ was detected with Peroxy Orange 1 (PO1;Tocris Bioscience 4944/10)). PO1 was dissolved in DMSO to make a 500 mM stock and was further diluted in water to make 10mM working solution. The stigmas were incubated in PO1 for 15 minutes in the dark and mounted in water for imaging. Mock treatments were done by incubating the stigmas in 0.1% DMSO for 15 minutes before imaging. Superoxide was detected using Dihydroethidium (DHE, Calbiochem 309800). DHE was diluted in Tris-HCL to make a 500 mM stock and was further diluted in water to make a working concentration of 10mM. The stigmas were incubated in DHE for 30 minutes in the dark and mounted in water before imaging. The images of stained stigmas were acquired on a Zeiss LSM 880 upright laser scanning confocal microscope using a W Plan-Apochromat 20x/0.8 M27 objective. PO1 and DHE were excited with a 488 nm & 514 nm laser at 0.25% power and emission was collected between 544-695 nm and 520-600 nm, respectively. The images were analyzed in Fiji. The mean fluorescence intensity of selected region of interest (ROI) was calculated in Fiji and the graphs were generated using GraphPad Prism version 10.3.1.

### NBT staining of stigmas for superoxide detection and quantification

Pistils from different stages (stage 9 – stage 13) were excised and vacuum infiltrated in Nitrotetrazolium blue (NBT; Sigma: MKCJ9460) stain (1mg/mL in 200 mM phosphate buffer pH 7.0 and 0.02% Silwet) and incubated in the staining solution for 90 minutes. Staining was performed on 5-10 pistils per stage. After staining, the samples were incubated in 70% ethanol and then cleared in (3:1:1) ethanol: glycerol: acetic acid at 98°C for 5 minutes. Samples were additionally cleared by mounting in chloral hydrate and visualized under a Nikon Ti2-E microscope equipped with DIC optics. Staining intensity quantification was performed using ImageJ software, version 1.52k (developed by Wayne Rasband, National Institutes of Health, Bethesda, Maryland, USA) and Java 1.8.0_172 (64-bit) engine. Images were converted into 8 bits and measured for image intensity after selecting the stained region of interest using the free hand selection tool to outline the region containing the papillae. Intensity numbers in the result window were converted into Optical Density (OD) numbers with the formula: OD= log (max intensity/Mean intensity), where max intensity = 255 for 8-bit images. Graphs were generated and statistical analysis was performed using Prism 9.0 software. OD value comparisons were made for individual developmental stages as well as pooled samples of all developmental stages of transgenic lines versus the wild-type.

### DAB staining of stigmas for hydrogen peroxide detection

Pistils from different stages (stage 9 – stage 13) from *Col-0* were excised and vacuum infiltrated with 3,3’-Diaminobenzidine (DAB; A.G. Scientific,Inc.:D-2561) stain (1 mg/ml in acidified 200 mM phosphate buffer and 0.02% silwet) and incubated in the staining solution for 4 hours. After staining, samples were stored in 70% ethanol and cleared in (3:1:1) ethanol: glycerol: acetic acid at 98°C for 5 min. Samples were additionally cleared by mounting in chloral hydrate and visualized under a Nikon Ti2-E microscope equipped with DIC optics.

### DPI and nPG treatments

Following emasculation of stage 12 flowers, the inflorescence stem was placed in 1 µM and 10 µM Diphenyleneiodonium chloride (DPI; Sigma: 43088) and 10 µM and 100 µM *n*-propyl gallate (*n*PG; ACROS organics:121-79-9) along with respective controls solutions and incubated overnight. Following treatments, pistils were collected and fixed in (9:1) ethanol: acetic acid overnight at 4º C. Samples were processed and observed under a Nikon Ti2-E microscope as described previously. Stigma papillae lengths were measured using ImageJ software, version 1.52k (developed by Wayne Rasband, National Institutes of Health, Bethesda, Maryland, USA) for 50 papillae from 3-4 flowers for each treatment. Prism 9 software (Graphpad) was used for plotting graphs and statistical analysis.

### Cloning and plant transformation

For the cloning of early expression stigma promoter *Chloris*, a region spanning 14,395 bp upstream of the START codon of AT5g53710 was amplified from *Arabidopsis thaliana* Columbia ecotype and introduced into Gateway vector pDONR207 using gene-specific primers with Gateway attB linkers (Supplementary Table S1). It was then recombined into pMDC163 GUS expression vector (Curtis and Grossniklaus, 2003) to produce in-frame pCHLORIS:GUS fusion construct that was then introduced into *Agrobacterium tumafaciens* strain GV3101 for plant transformation. Infusion cloning (Clontech) was used to introduce the pCHLORIS PCR fragment into the binary destination vector pGWB554 (addgene) after removing the CaMV35S promoter to enable gateway cloning (for primers, refer to Supplementary Table 1). The construct was verified by sequencing and designated as pSS13.16. Gateway cloning (LR reaction) was used to introduce AT3G10920 (*AtMSD1*) ORF entry clone cDNA in pENTR/SD-dTopo-vector (ABRC plasmid stock: U16034) into pSS13.16 to produce the *pCHLORIS:MSD1* fusion construct. A similar method was used to make the *pCHLORIS::CSD1/SOD1* (AT1G08830, ABRC plasmid stock: G11164) transformation plasmid. All constructs were transformed into *A. thaliana* Col-0 ecotype using the floral dip method (Clough and Bent, 1998). The transgenic plants were obtained by selecting the harvested seeds on MS medium (Murashige and Skoog, 1962) with 20 mg/L hygromycin, which were then transplanted to soil and grown in long days.

### Histochemical GUS assay and imaging

Flowers of different stages (stage 9 to stage 13) were collected from pCHLORIS:GUS T1 lines and fixed in 90% acetone for 40 min at -20 ºC. The samples were rinsed three times with phosphate buffer (100mM, pH 7.2) and vacuum infiltrated with GUS staining solution (0.1% Triton X-100, 10 mM EDTA, 2 mM ferrocyanide, 2 mM ferricyanide, 100 mM sodium dibasic phosphate, 100 mM sodium monobasic phosphate, and 4 mM 5-bromo-4-chloro-3-indoyl-β-D-glucuronide) for 5 min. The tissues were incubated at 37 ºC until GUS stain was visible. Stained flowers were incubated at room temperature overnight in 70% ethanol and stored at 4 ºC until imaging. Before imaging, the stained flowers were further cleared in 8:2:1 (w/v/v) chloral hydrate/ distilled water/glycerol solution. GUS staining pattern was whole flowers was imaged with an iPhone 8 camera and for stigma and anther specific expression it was visualized using a Nikon Ti2-E microscope equipped with DIC optics and verified in at least three independent lines to reduce uncertainty in expression patterns caused by positional effect of the insert.

### Expression analysis by RT-PCR

Unpollinated stage 12 stigmas from Col-0 and pCHLORIS:*MSD1* lines were collected in liquid nitrogen and stored in -80 ºC until analysis. Total RNA was extracted from these stigmas using E.Z.N.A Plant RNA kit (Omega BIO-TEK) according to manufacturer’s instructions. RNA was quantified using a NanoDrop spectrophotometer and RNA integrity was confirmed by resolving on an agarose gel. Total RNA was treated with DNAse I (Invitrogen) and reverse transcribed with 500 ng of RNA using superscript II reverse transcriptase (Invitrogen) according to manufacturer’s instruction. PCR amplification was performed with *MSD1* specific primers and *Ubq10* for internal reference (Supplementary Table S1). Each PCR reaction contained 10 µl 2X EmeraldAmp GT premix (Takara), 200 nM of each primer, and 1 µl of complementary DNA in final volume of 20 µl.

PCR amplification was performed for 28 cycles at 98 ºC for 10 sec, Annealing at 52 ºC for 30 sec and extension at 72 ºC for 60 sec with a preceding initial enzyme activation of 30 sec at 98 ºC. The PCR product was resolved in 0.8% Agarose gel and imaged with an AlphaImager (Alpha Innotech). Relative fold change in transcript levels were calculated by quantifying band intensities in the Agarose gel using ImageJ.

## Acknowledgements

We thank Dr. Yan Ju and Sienna Ogawa for critical reading of the manuscript, the Kessler lab, Dr. Leonor Boavida, Dr. Daniel Szymanski and Dr. Yun Zhou for helpful discussions. We thank Dr. Thierry Gaude and Isabelle Fobis-Loisy for the CMT marker line. This work was funded by startup funds from Purdue University to S.A.K.

## Author Contributions

All authors conceived the experimental questions and approach; S.S., S.D.V., and T.C.D. designed and performed experiments; all authors analyzed the data and wrote the manuscript.

## Supplemental Figure Legends

**Figure S1.**
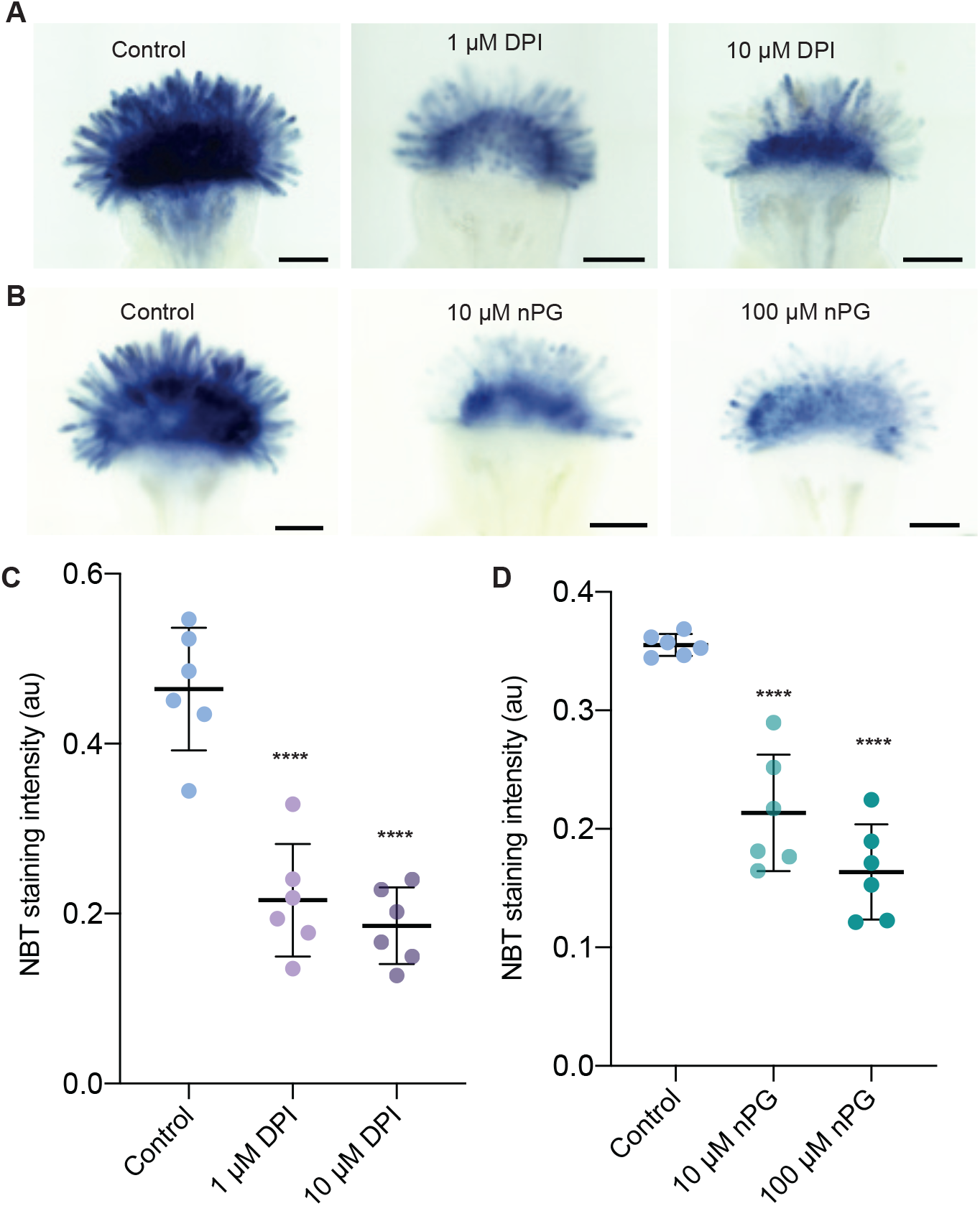
Superoxide levels are reduced following transpirational uptake of DPI and nPG by the inflorescence. (A) Representative image of NBT stained stigmas of control and DPI (1 µM and 10 µM) transpirational uptake. (B) Average NBT stain intensity of stigmas from control and following DPI transpirational uptake (n=6). (C) Representative image of NBT stained stigmas of control and following nPG transpirational uptake (10 µM and 100 µM). (D) Average NBT stain intensity of stigmas from control and following nPG transpirational uptake (n=6) Error bar indicate ± SD. Asterisks indicate significant differences (****P<0.0001) determined by Dunnett’s test. Scale bars=100 µm.

**Figure S2.**
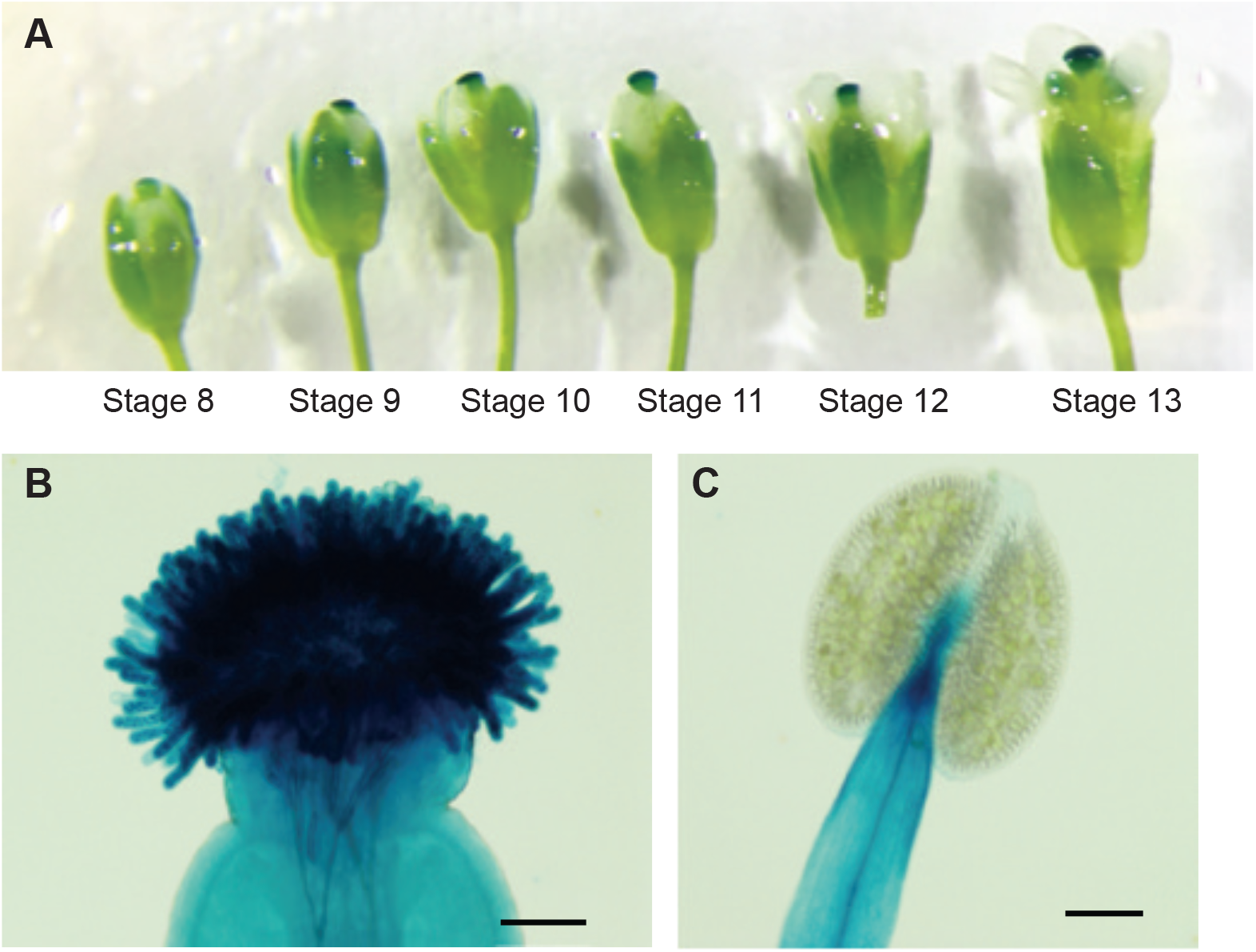
The CHLORIS promoter is active throughout stigma development and not expressed in pollen. (A) pCHLORIS::GUS expression (blue) at different stages of flower development as visualized by GUS assay. (B) pCHLORIS::GUS expression in a mature stigma. (C) pCHLORIS::GUS expression in a mature stamen. Scale bars = 100 µm.

**Figure S3.**
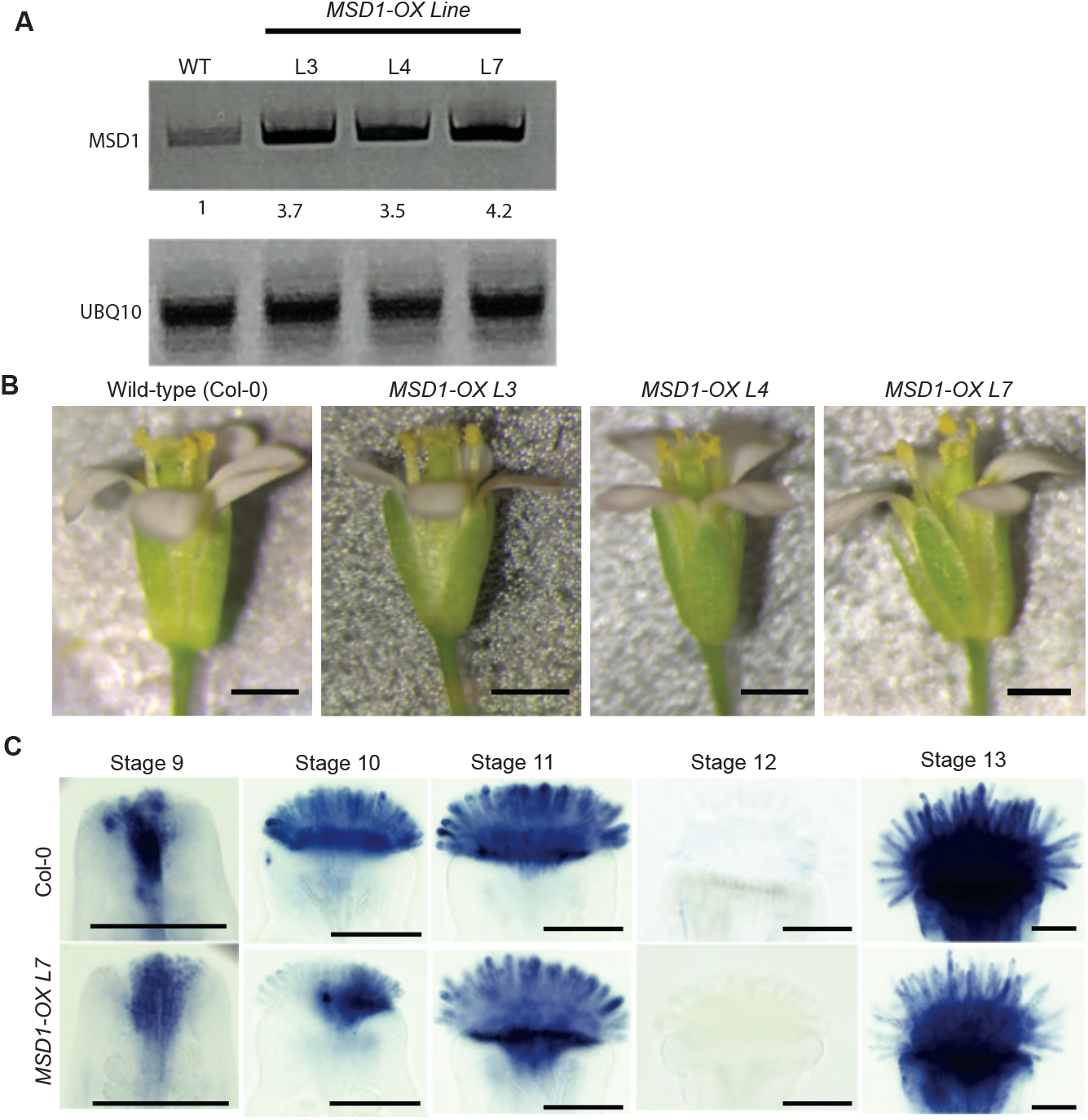
Genetic evidence for superoxide’s role in stigma papillae development. (A) RT-PCR analysis showing enhanced *MSD1* transcript levels in pCHLORIS::MSD1 lines (numbers under MSD1 panel indicate fold change compared to Col-0). (B) Flowers from wild-type (Col-0) and three *MSD1-OX* lines. (C) NBT stained stigmas of wild-type and MSD1 over-expression line (L7) at various stages of flower development. Scale bars = 100 µm.

## Notes

### Competing Interest Statement

The authors have declared no competing interest.

